# Antibiotic Stress Selects against Cooperation in a Pathogenic Bacterium

**DOI:** 10.1101/053066

**Authors:** Marie Vasse, Robert Noble, Andrei R. Akhmetzhanov, Clara Torres-Barceló, James Gurney, Simon Benateau, Claire Gougat-Barbera, Oliver Kaltz, Michael E. Hochberg

**Affiliations:** Institut des Sciences de l’Evolution, CNRS-Université de Montpellier, Place Eugène Bataillon, Montpellier Cedex 5 34095, France; Institute of Statistical Science, Academia Sinica, 128 Academia Rd, Taipei, Taiwan; Santa Fe Institute, 1399 Hyde Park Road, Santa Fe, NM 87501, USA

## Abstract

Ecological antagonisms such as predation, parasitism, competition, and abiotic environmental stress play key roles in shaping population biology, in particular by inducing stress responses and selecting for tolerant or resistant phenotypes. Little is known, however, about their impact on social traits, such as the production of public goods. Evolutionary trade-off theory predicts that adaptation to stresses should lessen investments in costly helping behaviours when cooperation does not increase resistance or tolerance, but support for this prediction is scarce. We employed theory and experiments to investigate how ecological antagonism influences social dynamics and resistance evolution in the pathogenic bacterium *Pseudomonas aeruginosa*. We subjected two clones of bacterium to four doses of antibiotics and assessed growth and frequencies of public goods producing and non-producing genotypes. Our results show that abiotic stress selects against public goods production. Specifically, we found that non-producers of costly iron chelating molecules (siderophores) most rapidly increased in frequency under intermediate antibiotic pressure. Moreover, the dominance of non-producers in mixed cultures was associated with higher survival and resistance to antibiotics than in either producer or non-producer monocultures. Mathematical modelling explains this counterintuitive result, and shows how these qualitative patterns are predicted to generalise to many other systems. Our results shed light on the complex interactions between social traits and ecological antagonisms, and in particular the consequences for bacterial social evolution and antibiotic resistance.

## Introduction

Public goods production is a characteristic of a diverse range of taxa, from microbes to humans (1–3). Explaining the persistence of this costly behaviour is challenging, since cheats can exploit the commons without contributing. Kin selection theory is a successful framework for understanding the evolutionary dynamics of public goods behaviours, with the central prediction that cooperation is favoured by sufficient benefits to, and positive assortment between, cooperators (4–8).

Microbial populations are an increasing focus for understanding public goods dynamics (9–11). Microbes may exhibit rapid ecological and evolutionary responses and are amenable to controlled laboratory experimentation (10,12). Bacteria, in particular, show a variety of behaviours consistent with basic social interactions. These frequently involve the coordinated secretion of metabolites that are potentially beneficial to others (i.e., public goods), leading to, for example, collective motility and/or resource acquisition (reviewed by e.g., (9,13,14)).

Recent study in experimental bacterial populations has elucidated mechanisms such as assortment emerging from limited or budding dispersal (15,16) and kin discrimination (17,18) that are consistent with kin selection fostering cooperative behaviours (e.g., (8,19)). Despite this accumulating consensus, little is known about how social populations respond to differences and variation in abiotic and biotic components of their environment, and in particular, ecological antagonisms. Under the assumption that stress responses (20–22) or the evolution of stress resistance (23–25) entail costs, they could potentially interact with social behaviours and their evolution.

Bacteria are confronted with a variety of antagonisms, including predation and parasitism (e.g., phages, metazoans, plasmids), antimicrobials produced by other organisms (antibiotics, AMPs, toxins), and abiotic environments (extreme temperatures, pH, salinity) that can result in reduced fitness through decreases in survival and reproduction. In particular, subinhibitory concentrations of antimicrobials are pervasive in natural environments such as rivers, lakes, soils, and bacterial hosts (animals and plants) and play a key role in progressive increases in antibiotic resistance (26–28). Such concentrations were shown to affect cellular physiology, genetic variability and behaviours with mostly unknown evolutionary implications (26). Social behaviour, namely cooperation, may be affected by stress directly through differential selection on cooperative phenotypes (29,30), or by inducing specific plastic behaviours (31–35), in particular when cooperation leads to increased resistance or tolerance to stress. In the absence of a direct benefit of the cooperative behaviour against stress, cooperation may nonetheless be influenced indirectly through impacts on population structure and dynamics (36), via epistasis or pleiotropy (37), or through the hitchhiking of cooperative genes with resistance mutations (38–40).

If cooperation does not have a direct benefit (35), the ecological and evolutionary outcomes of the interactions between non-defensive public goods and stress responses are potentially complex. In principle, because the cooperative trait is not under direct selection, its spread in the population might rely on hitchhiking with resistant or tolerant genotypes (38,40,41). As the probability of the emergence of resistance or tolerance mutations should increase with population size (38,42), such mutations are most likely to appear in the more numerous subpopulation and/or in the most competitive subpopulation (i.e., with the fastest growth). This would suggest that the fate of cooperators is in part contingent on whether they represent the majority of the population when the environmental stress occurs. More generally, however, we would expect that, under sublethal stress, non-producers limit the emergence and spread of resistant cooperators by cheating on public-goods production.

The above considerations lead to the prediction that the direct cost of public goods production and indirect costs through exploitation by non-producers accentuate both the ecological and evolutionary benefits of cheating when the population faces an environmental stress. We experimentally tested this prediction by examining how a public goods trait in the form of siderophore production interacts with resistance evolution to the antibiotic gentamicin in the pathogenic bacterium *Pseudomonas aeruginosa*. We grew a strain of *P. aeruginosa* PAO1 producing the siderophore pyoverdin and/or a non-producing strain under iron-limited conditions with different doses of the antibiotic gentamicin. Siderophores are small secreted molecules that chelate poorly-soluble iron in the environment, making the iron available to bacteria via specific outer-membrane receptors (43). As any cell carrying these receptors can use chelated iron, siderophores are a public good in well-mixed environments. As such, costly siderophore production is vulnerable to ‘cheating’ by cells that do not produce, but possess specialized receptors and reap the benefit of available iron (e.g., (19,44)). We used subinhibitory antibiotic concentrations, which the bacteria are most likely to encounter in natural settings and in host tissues (45–48). We assessed (i) the impact of antibiotic pressure on the interaction between the two production genotypes and (ii) the consequences for the population response to antibiotics, in particular the evolution of resistance. We found that antibiotic stress accelerates the decline of producers in mixed cultures. Moreover, non-producer resistance frequency was greater in mixed cultures compared to monocultures of either non-producers or producers. A mathematical model shows that these observed qualitative patterns may be explained by the constitutive investment in pyoverdin production decreasing the capacity of producers to cope with antibiotic stress in the presence of non-producers. Given the generality of the model, its predictions regarding the interplay between social and resistance traits should apply to many other biological systems. We discuss these findings in the contexts of social evolution and resistance to antagonisms, with a focus on bacterial evolution.

## Experimental results

### 1. Changes in non-producer frequency

In all mixed populations, for all three initial frequencies and four antibiotic treatments (Fig 1A), non-producer frequency increased over the course of the experiment (Fig 1B). Non-producer frequencies were substantially higher in the presence of the antibiotic than in the antibiotic-free controls. At higher doses (4 and 8 μg/mL), non-producers had often reached near-fixation (> 90%) after 48 hours.

**Fig. 1:**
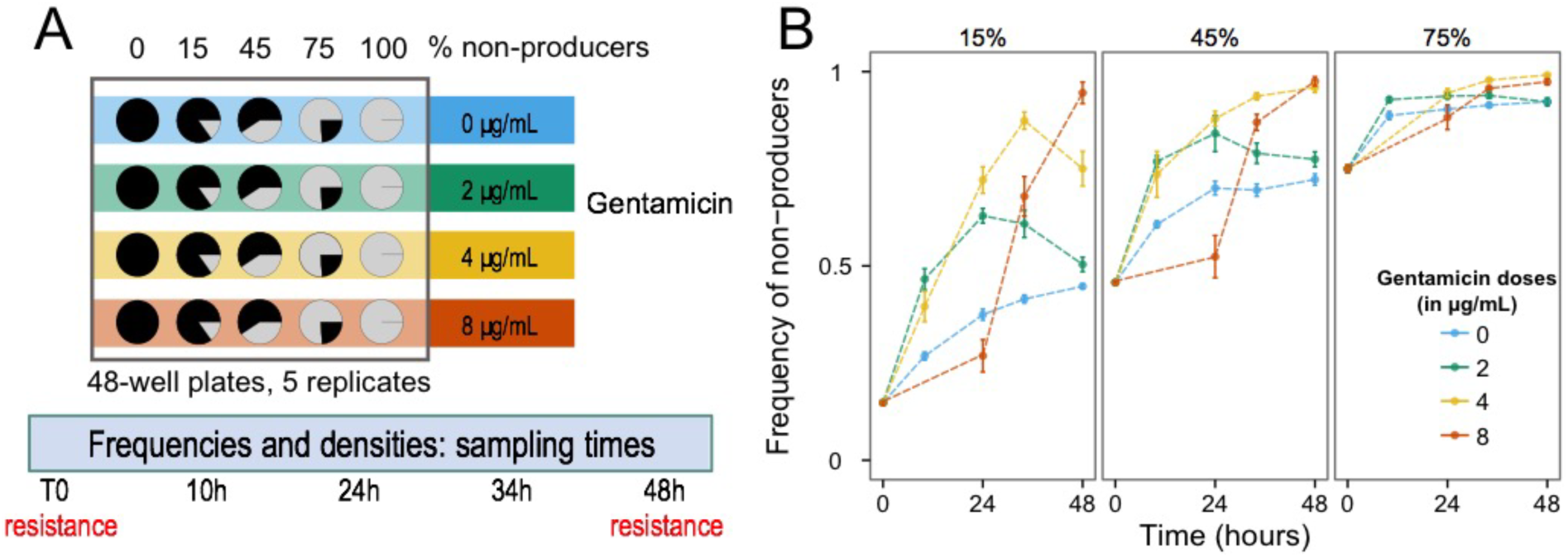
Experimental design (A) and change in non-producer frequencies in experimental populations between beginning (T0) and end of the experiment (T48) (B). (A) Experimental design. (B) The three panels correspond to the different initial frequencies of non-producers. Colours represent gentamicin doses: blue = 0 μg/mL, green = 2 μg/mL, yellow = 4 μg/mL and red = 8 μg/mL. Bars are standard errors of the mean.

Antibiotic dose further affected the timing of changes in the relative frequencies of producers and non-producers, as indicated by the significant time × gentamicin interaction (*χ*^*2*^_3_ = 123.39, *ρ* < 0.0001). Namely, at the two lower doses (2 and 4 μg/mL), non-producer frequencies increased considerably during the first 24 hours and then reached a peak (Fig 1B). At the highest antibiotic dose (8 μg/mL), this increase was delayed by *c* 24h.

These patterns were similar for all initial frequencies of non-producers, and the significant three-way interaction (time × gentamicin × initial frequency, *χ*^*2*^_*6*_ = 26.69, *ρ* < 0.001) likely reflects the lower absolute change in frequency for populations initiated with 75% of non-producers (since frequencies cannot exceed 100%).

### 2. Effects of the antibiotic and initial non-producer frequency on bacterial antibiotic resistance

Experimental treatments affected resistance frequencies in three main ways. First, increasing the dose of gentamicin led to higher frequencies of resistant cells, with up to a five-order-of-magnitude difference between the highest dose (8 μg/mL) and the control (Fig 2). Nonetheless, the frequency of resistance always remained below 10%.

**Fig. 2:.**
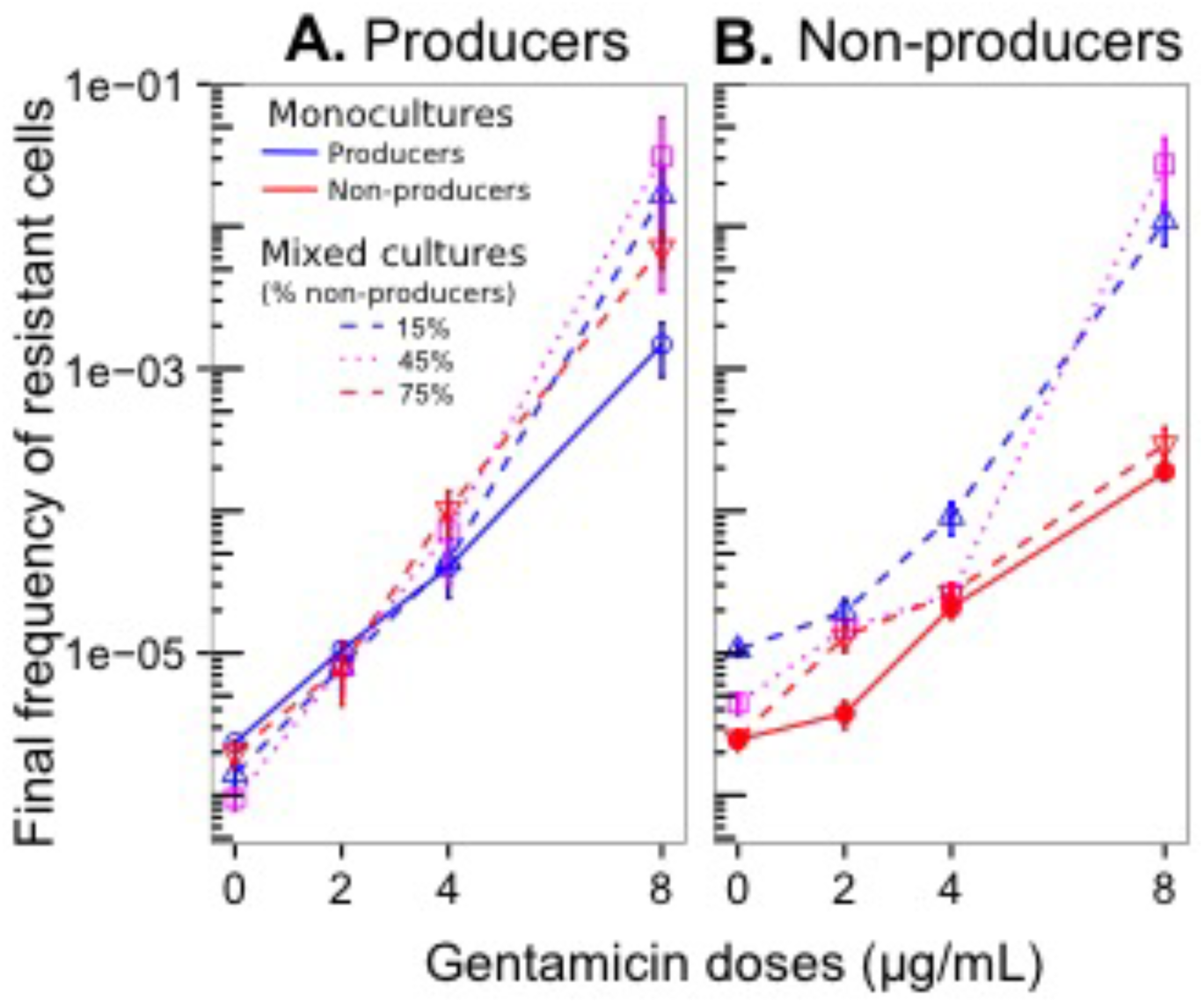
Antibiotic resistance in experimental populations at end of the experiment (T48). Final frequency of esistant cells in producers (A) and in non-producers (B) in monocultures (solid lines) and in mixed cultures (dashed lines) for different doses of gentamicin. Bars are standard errors of the mean.

Second, producer monocultures generally showed higher frequencies of resistance than non-producer monocultures at all three gentamicin doses (producer vs. non-producer: *χ*^*2*^_*1*_ = 9.0, *ρ* = 0.0027; Fig 2). Third, resistance was more frequent in mixed culture than in monoculture, in particular at high antibiotic dose (Fig 2). This general pattern held for both producers (mono vs. mixed: *χ*^*2*^_*1*_ = 43.7, *ρ* < 0.0001) and non-producers (*χ*^*2*^_*1*_ = 34.2, *ρ* < 0.0001), despite some deviations (significant dose × initial non-producer frequency interactions for both producer types: *χ*^*2*^_*9*_ > 16, *ρ* < 0.05). Namely, for producers, the higher mixed-culture resistance was only consistently prominent in lines from the highest antibiotic dose treatment (Fig 2A). Non-producers showed higher mixed-culture resistance over all dose treatments, but there was more variation among lines with different initial non-producer frequencies (Fig 2B). A supplementary replicate experiment confirmed these main results (Fig S5).

The unexpected observation of higher frequencies of producer resistance in mixed culture led us to conduct a series of additional assays (Appendix S2) to investigate in more detail the quantitative levels of resistance (measured as the minimum inhibitory concentration) and associated fitness costs in mixed and monocultures. Our hypothesis was that producers in mixed culture might have acquired particular adaptations to selection pressures from both non-producers and the antibiotic, resulting in highly resistant, fit producers that could coexist with non-producers. Indeed, we found that, unlike non-producers, producers tended to be more resistant (higher level of resistance) in mixed cultures compared to monocultures (*ρ* < 0.05, Fig S6) and this higher resistance did not come at an increased fitness cost (Fig S7). Based on the growth assays, we did not detect significant differences in pyoverdin availability (Fig S9) nor production per cell (Fig S10) between resistant colonies from mixed cultures and from monocultures. This suggests that competition with non-producers did not select for decreased pyoverdin production in resistant producers.

We then investigated genetic resistance to gentamicin in resistant and non-resistant individual colonies of producers and non-producers from all treatments in the repeated 48-hour experiment (Appendix S2). We sequenced the repressor gene and intergenic region of the efflux pump MexXY, described as the only identified pump mediating aminoglycoside resistance (49,50). While the observed selection for resistant phenotypes suggests an underlying genetic component, we did not detect any evidence of gene modification in these markers (Appendix S2). We further tested for the presence of 9 genes coding for gentamicin-degrading enzymes and did not detect any synonymous nor non-synonymous modifications of these genes.

## Theoretical framework

Our experimental results showed that (i) antibiotics increased the frequency of non-producers in mixed cultures in a dose-dependent manner and (ii) the frequency of resistant cells was higher in mixed cultures than in either monoculture. We hypothesised that the cost of pyoverdin production reduced the capacity of producers to cope with antibiotic stress, perhaps by depleting metabolic resources that would otherwise be expended on countering the drug’s effects. This would be especially pronounced in the presence of non-producers since the latter constitute an indirect cost by removing iron from the environment. We developed and analysed a mathematical model to examine this hypothesis and to investigate the interactions between the effects of antibiotics and pyoverdin.

### 1. Effects of antibiotics on the interactions between producers and non-producers

To test the generality of our findings, we constructed a dynamical model of a population of producers and non-producers, each composed of antibiotic-resistant and susceptible subpopulations (Appendix S4). We assumed the fitness of each subpopulation depends on the cost of pyoverdin production, the cost of antibiotic resistance, the population density, the beneficial effect of pyoverdin, and effects of the antibiotic on bacterial growth. Analysis of this model shows that non-producer frequency is expected to increase fastest at intermediate antibiotic concentrations if the antibiotic (i) affects producers more than non-producers at low and intermediate concentrations, but (ii) impacts both populations similarly at very high concentrations. The pattern holds for a wide range of plausible parameter values, and is not sensitive to pyoverdin dynamics (Appendix S4).

To further quantify antibiotic effects in our experimental system, we tailored the mathematical model to our particular biological system by specifying functional forms for each of the factors contributing to bacterial fitness (Appendix S4). The resulting dynamical model provides a good statistical fit to the experimental data (Fig S15) and shows frequency dynamics qualitatively consistent with experimental observations (Fig 3). Initially, non-producer frequency increases fastest at intermediate antibiotic concentrations, and increases slowest at high antibiotic concentrations. However, under the reasonable assumption that the antibiotic effect decreases over time, the model predicts that the rate of change will accelerate in the latter case, as we indeed observed in our experiments.

**Fig. 3:**
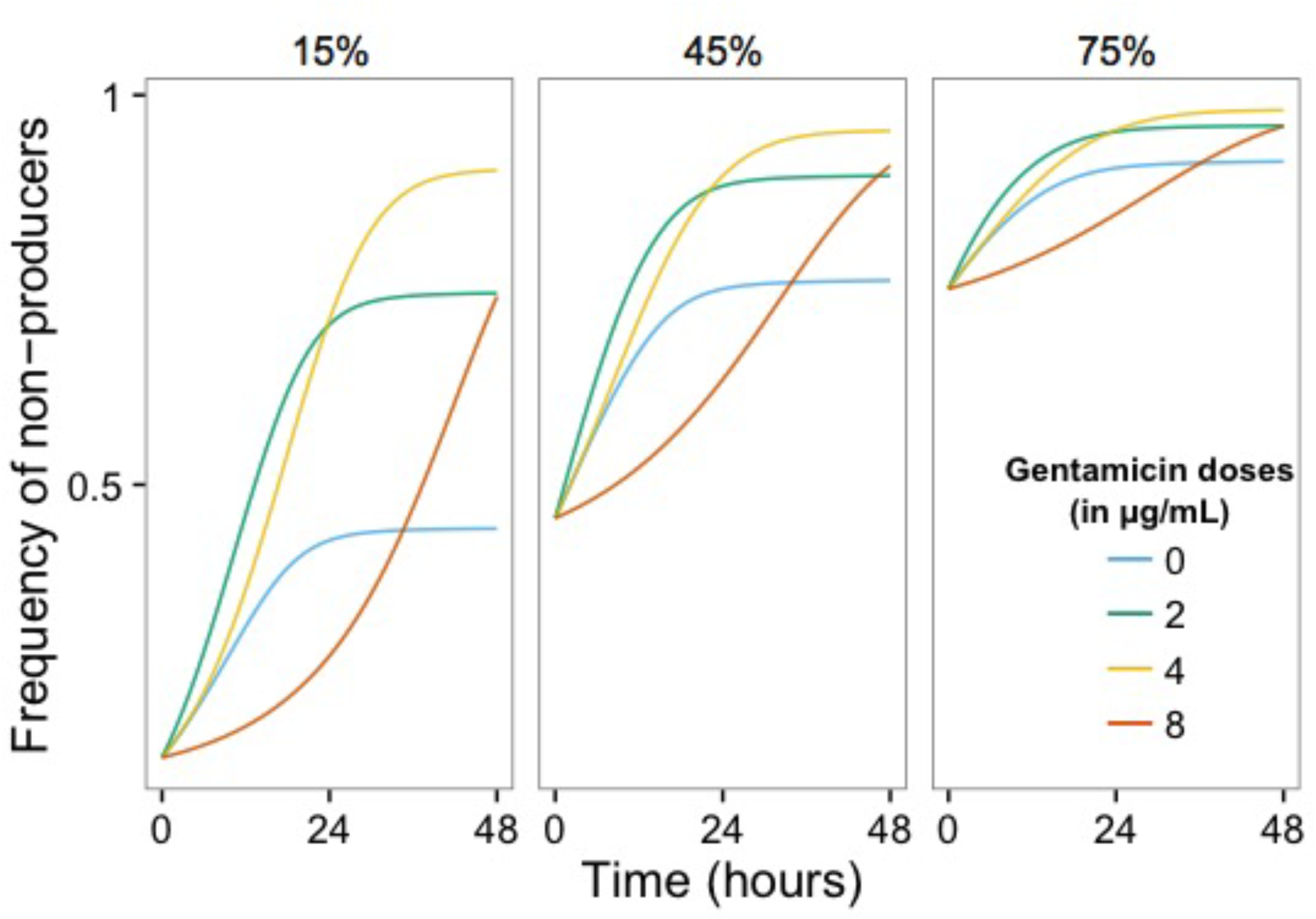
Change in non-producer frequencies in a mathematical model between T0 and T48. Mathematical model behaviour based on statistical parameter estimations from experimental data (*cf*. Fig 1B for experimental data). The three panels correspond to the different initial frequencies of non-producers. Colours represent gentamicin doses: blue = 0 μg/mL, green = 2 μg/mL, yellow = 4 μg/mL and red = 8 μg/mL. Model definition and parameters values are in Appendix S4.

Based on parameter values of the fitted model (Appendix S4, Table S4), an antibiotic concentration of 2 μg/mL reduces the growth rate of producers by approximately 55% and the growth rate of non-producers by only 19% in mixed cultures, which leads to a rapid increase in non-producer frequency. At 8 μg/mL, the growth rates are reduced by approximately 99% and 93% respectively, resulting in a much slower increase in non-producer frequency. The model indicates that the antibiotic effect declines approximately 60% over the course of each experiment.

### 2. Effects of antibiotic dose and initial non-producer frequency on antibiotic resistance

We next investigated the effects of the initial frequency of non-producers (and antibiotic dose) on the frequency of resistant cells in evolved populations. Analysis of our mathematical model (Appendix S4) shows that, for high antibiotic doses, a public good (such as pyoverdin) will increase the final frequency of antibiotic-resistance if (i) the beneficial effect of the public good interacts with the antibiotic effect or with the cost of resistance (or both); and (ii) the antibiotic is primarily bacteriostatic. When these conditions are met, the beneficial effects of the public good accentuate the fitness difference that results from the unequal effects of the drug on susceptible and resistant bacteria. By fitting the model to our data (Appendix S4, Fig S15), we found that the size of this interaction effect increases with antibiotic dose within the range tested in our experiments (Fig 4).

**Fig. 4:**
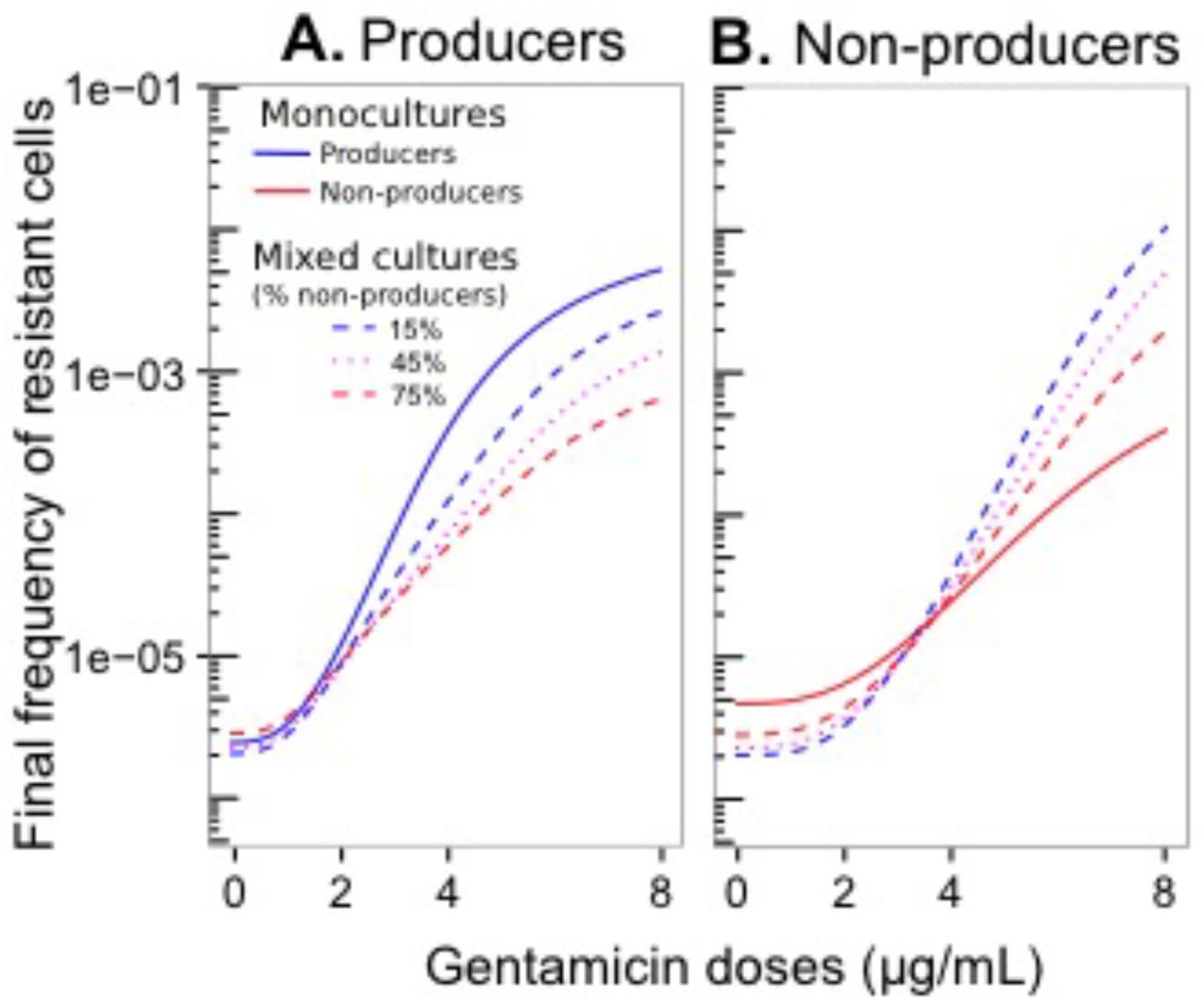
Resistance frequencies in a mathematical model at T48. Mathematical model behaviour based on statistical parameter estimations from bacterial population data (*cf.* Fig 2 for experimental data). Final frequency of resistant cells in producers (A) and non-producers (B) in monocultures (solid lines) and in mixed cultures (dashed lines) for different doses of gentamicin. Model definition and parameters values are in Appendix S4.

## Discussion

Social behaviours are widespread across the living world at all organisational levels (51). While underlying intrapopulation interactions have been extensively studied, their interplay with environmental factors remains poorly understood. Bacteria represent an excellent model system for exploring these processes in large populations under controlled conditions (13). Here, we focused on the interplay between antibiotic stress and siderophore cooperation in *P. aeruginosa*. We found that antibiotics constitute a cost to social behaviours, manifested by an increased benefit of cheating under stressful conditions. Mathematical analyses show the key driver to be the differential stress sensitivities of the two public goods strategies. Our experimental and theoretical results contribute to disentangling the complex ecological and evolutionary dynamics of public goods behaviours and their interactions with biological stressors such as antibiotics. Below we discuss the importance of ecological antagonisms in the evolution of public goods behaviours and resistance.

The essential element underlying all of our results is that, while public goods benefit every cell, only producers pay the associated fitness cost. This factor can explain why producers were more affected than non-producers by antibiotic stress: the fitness cost of pyoverdin production limited the capacity of producers to resist antibiotics and to compete with non-producers. In other words, in the absence of a ‘private benefit’ to producers (which would be expected to operate, for example, in spatially structured environments), it is growth in the absence of antibiotics that, all else being equal, determines how well a strain can cope with an ecological antagonism. This is consistent with the findings of Mitri and collaborators (52) who employed computer simulations in a spatial setting and showed that antagonism (in this case ecological competition) is more detrimental to cooperators than to cheats when nutrients are limiting. The authors suggested that this effect is due to the investment in cooperative secretions limiting growth and thereby competitive ability. Alternatively, in some particular cases, antagonisms may directly increase the benefit to non-producers by inducing the production of costly public goods (35,53). In *Staphylococcus aureus*, for example, sublethal doses of ciprofloxacin, mupirocin, or rifampicin induce the expression of the costly effector molecule regulating the quorum-sensing system *agr*, thereby favouring *agr*-negative variants (53).

In one of the few studies investigating interactions between antibiotics and social behaviours in bacteria, Diard and colleagues (29) assessed the *in vivo* impact of ciprofloxacin on competition between virulent cooperative *Salmonella enterica* serovar Typhimurium and avirulent defectors. In the absence of the antibiotic, defectors outcompeted cooperators in the gut lumen, whereas the antibiotic addition reversed the outcome, leading to selection for the virulent cooperators. Indeed, only the virulent cells were able to invade host tissues and escape antibiotic mortality in the lumen. The authors observed that, when antibiotic pressure decreased, the virulent strain reinvaded the gut lumen. Our results contrast with these findings of Diard and coworkers, as we observed that antibiotics led to the selection of non-producers over producers. We hypothesize that this is because of differences in spatial structure between the two studies: whereas the gut lumen is a highly structured environment, our *in vitro* studies were conducted under well-mixed conditions, where producers had no refuge from the antibiotic, nor from exploitation by non-producers.

Whereas pyoverdin availability had relatively little effect on non-producer frequency dynamics, it had a major role in resistance evolution. Producer monocultures should have the highest per-cell pyoverdin availability, leading to the highest growth rates and population sizes, and therefore one might expect to see the highest final frequency of resistance in producer monocultures. However, we found that, under the highest antibiotic dose, the frequency of resistant cells was higher in mixed cultures. Our mathematical model shows that this pattern is predicted when a bacteriostatic antibiotic affects producers more than non-producers, provided the beneficial effect of the public good interacts with the effects of the drug (Appendix S4). This can explain why resistant non-producers grew faster in mixed cultures, not only compared to resistant non-producers in monoculture, but also compared to resistant producers in monoculture, thereby leading to more resistance in mixed populations. The optimal frequency for non-producers appears to be low. When initially very frequent (75%), non-producers did not evolve substantially higher frequencies of resistance compared to their populations in monoculture, possibly as uncommon producers yielded low pyoverdin concentration thereby contributing relatively little both to non-producer growth (35) and to the generation of resistant variants. An additional experiment (Appendix S3) confirmed that differences in growth and antibiotic resistance between producers and non-producers are mediated by pyoverdin availability and production: when populations grew under high iron availability conditions (i.e., where siderophores are not needed), the frequency of resistance was similar for both strains in mixed cultures and in both monocultures (Fig S13). Our mathematical model also predicts that, when iron availability is limited, resistance among producers will be more frequent in monocultures than in mixed cultures, yet experimentally we observed the opposite pattern. This discrepancy between theory and experiments may be explained by the steep decline in sensitive producer densities near the end of the culture period, as they succumbed to the combined effects of antibiotics and exploitation by cheats.

Our results have implications for the control of pathogenic bacteria using antibiotics, and especially for the treatment of multiple versus single strain infections, as we have found that antibiotics can select for an overall higher prevalence of resistance when both producers and non-producers are present. It has previously been shown that selection for non-producers is expected to lower bacterial virulence (54,55). We observed that highly resistant producers also arose in mixed cultures, but they were outcompeted by resistant non-producers. This raises the question of how the two types can coexist (resulting in more virulent infections) as has been observed *in situ* (56). A reasonable hypothesis is spatial structure (57–60), whereby spatial segregation of non-producers and producers limits the exploitative potential of the former. Future theoretical and experimental study is needed to test this prediction.

Overall, our results indicate that ecological stressors can play an important role in public goods dynamics by affecting the interactions between producers and non-producers. In turn, these interactions feed back to the population’s ecological and evolutionary responses to the stressors, affecting survival and resistance. Previous research has propounded the importance of ecology in social evolution and called for a deeper integration of ecological factors in social theory (61–65). Further to this claim, we suggest that ecological stressors could impact social evolution in microbes and in multicellular taxa more generally. Testing this broader hypothesis would require extensions of our mathematical model and experiments to assess the costs and payoffs of different social strategies in a wider range of environments with various types of spatial structure.

## Materials and Methods

### Experiment

#### Experimental protocol

We used *P. aeruginosa* PAO1 as the pyoverdin producing wild type (‘producers’) and an otherwise isogenic mutant PAO1Δ*pvdD* (66) unable to produce pyoverdin (‘non-producers’). We inoculated bacteria as either monocultures or mixed cultures of producers and non-producers to a final density of *c* 10^7^ bacteria per mL into fresh iron-limited medium with either a low (2 μg/mL), intermediate (4 μg/mL), or high (8 μg/mL) dose of gentamicin, or in antibiotic-free medium (Fig 1A). Mixed populations were initiated with 15%, or 45%, or 75% of non-producers. Each treatment was replicated 5 times for a total of 100 populations (4 antibiotic conditions × 5 types of cultures × 5 replicates) that were arbitrarily distributed in the 48-well plates. The experiment was run for 48 hours at 37°C under constant shaking (350 rpm, 8 mm stroke). We measured the densities and the relative frequencies of producers and non-producers by plating samples of each population onto King’s B medium (KB) agar plates and subsequent counting of colony forming units (CFUs) at the beginning of the experiment (T0) and after 10 (T10), 24 (T24), 34 (T34) and 48 (T48) hours. In addition, we estimated the frequency of resistant cells at T0 and T48 by plating samples of each population onto antibiotic-free KB agar plates and onto KB agar with 10 μg/mL gentamicin, simultaneously. Experimental conditions are detailed in Supplementary materials (Appendix S1).

#### Statistical analysis

We employed logistic regression (binomial error structure, logit link) to analyse variation in the frequency of non-producer cells over the course of the experiment. Models contained antibiotic treatment (0, 2, 4, 8 μg/ml gentamicin) and initial non-producer frequency (0, 15, 45, 75 and 100%) as explanatory factors, time (hours) as a covariate, and replicate population as a random factor. All possible interactions were statistically fitted. Where necessary in this and other analyses below, models were corrected for overdispersion by dividing the *χ*^*2*^ values for each factor by the mean residual deviance (*χ*^*2*^ value of residuals divided by the degrees of freedom of residuals).

We analysed variation in the frequency of antibiotic-resistant cells as a function of antibiotic treatment and initial non-producer frequency. Attempts using logistic regression, with the number of resistant cells as the response variable and total cell number as the binomial denominator (as above for non-producer frequency, with logit link) led to unsatisfactory model fits, namely a strong deviation from a normal distribution of residuals. This problem was solved by switching to a Poisson error structure (with log link) and using total cell number as a covariate. This method is also more appropriate, given that numbers of resistant and non-resistant cells were determined independently (by plating independent samples on different media, see above). Analyses were carried out separately for producers and non-producers. We employed orthogonal contrasts for comparisons between mono-and mixed culture treatments.

#### Mathematical analysis and modelling

To explore the generality of our experimental results, we conducted a mathematical analysis of the relative fitness of producer and non-producer bacterial populations, and of antibiotic resistant and susceptible subpopulations. The main assumptions underlying this analysis are that pyoverdin production and antibiotic resistance incur fitness costs, and the antibiotic effects on producers and non-producers may be unequal. The full analysis can be found in Supplementary materials (Appendix S4).

We also developed a dynamical mathematical model as a quantitative test of our assumptions. The model assumes that the cost of pyoverdin production is proportional to the production rate, which is inversely related to pyoverdin concentration. Pyoverdin is assumed to increase both bacterial fission rate and carrying capacity. The primary effect of the antibiotic is bacteriostatic, and is a sigmoidal function of antibiotic concentration. We further assumed that the antibiotic effect decreases over time, which could occur, for example, due to drug degradation linked to the accumulation of protective compounds (such as degrading enzymes, e.g., (67) or from bacterial adaptation (68). We fitted the model to the bacterial population data using a Markov chain Monte Carlo (MCMC) method (69,70) and verified the fit with a different algorithm (71). Further details of the model are in Supplementary materials (Appendix S4).

## Acknowledgements

We thank Pierre Cornelis and Melanie Ghoul for providing the PAO1 and PAO1Δ*pvdD* strains of *P. aeruginosa*; David Lunn for helping us acquire and set up software; and Guillaume Martin, Sylvain Gandon, Sonia Kéfi, Sébastien Lion and Rolf Kümmerli for helpful advice.

## Funding

This work was supported by James S. McDonnell Foundation Studying Complex Systems Research Award No. 220020294 (to M.E.H.) and a doctoral grant from the French Ministry of Research (to M.V). The funders had no role in study design, data collection and analysis, decision to publish, or preparation of the manuscript.

## Competing interests

We declare we have no competing interests.

